# A Network of Networks Approach for Modeling Interconnected Brain Tissue-Specific Networks

**DOI:** 10.1101/349969

**Authors:** Hideko Kawakubo, Yusuke Matsui, Itaru Kushima, Norio Ozaki, Teppei Shimamura

## Abstract

**Motivation:** Recent sequence-based analyses have identified a lot of gene variants that may contribute to neurogenetic disorders such as autism spectrum disorder and schizophrenia. Several state-of-the-art network-based analyses have been proposed for mechanical understanding of genetic variants in neurogenetic disorders. However, these methods were mainly designed for modeling and analyzing single networks that do not interact with or depend on other networks, and thus cannot capture the properties between interdependent systems in brain-specific tissues, circuits, and regions which are connected each other and affect behavior and cognitive processes.

**Results:** We introduce a novel and efficient framework, called a “Network of Networks” (NoN) approach, to infer the interconnectivity structure between multiple networks where the response and the predictor variables are topological information matrices of given networks. We also propose Graph-Oriented SParsE Learning (GOSPEL), a new sparse structural learning algorithm for network graph data to identify a subset of the topological information matrices of the predictors related to the response. We demonstrate on simulated data that GOSPEL outperforms existing kernel-based algorithms in terms of F-measure. On real data from human brain region-specific functional networks associated with the autism risk genes, we show that the NoN model provides insights on the autism-associated interconnectivity structure between functional interaction networks and a comprehensive understanding of the genetic basis of autism across diverse regions of the brain.

**Availability:** Our software is available from https://github.com/infinite-point/GOSPEL.

**Supplementary information:** Supplementary data are available at *Bioinformatics* online.

## 1 Introduction

Neurodevelopmental disorders are characterized by impaired functions of the central nervous system that can appear early in development and often persist into adulthood (Tollefsbol, 2017). The spectrum of developmental impairment varies and includes intellectual disabilities,communication and social interaction challenges, and attention and executive function deficits (American Psychiatric Association, 2013).Prototypical examples of neurodevelopmental disorders are intellectual disability, autism spectrum disorder (ASD), epilepsy, and schizophrenia.

Recent sequence-based analyses have unraveled a complex, polygenic,and pleiotropic genetic architecture of neurodevelopmental disorders, and have identified valuable catalogs of genetic variants as genetic risk factors for neurodevelopmental disorders (Gratten *et al.*, 2014). However, it remains unknown if and how genetic variants interact with environmental and epigenetic risk factors to impart brain dysfunction or pathology.

**Fig. 1.**
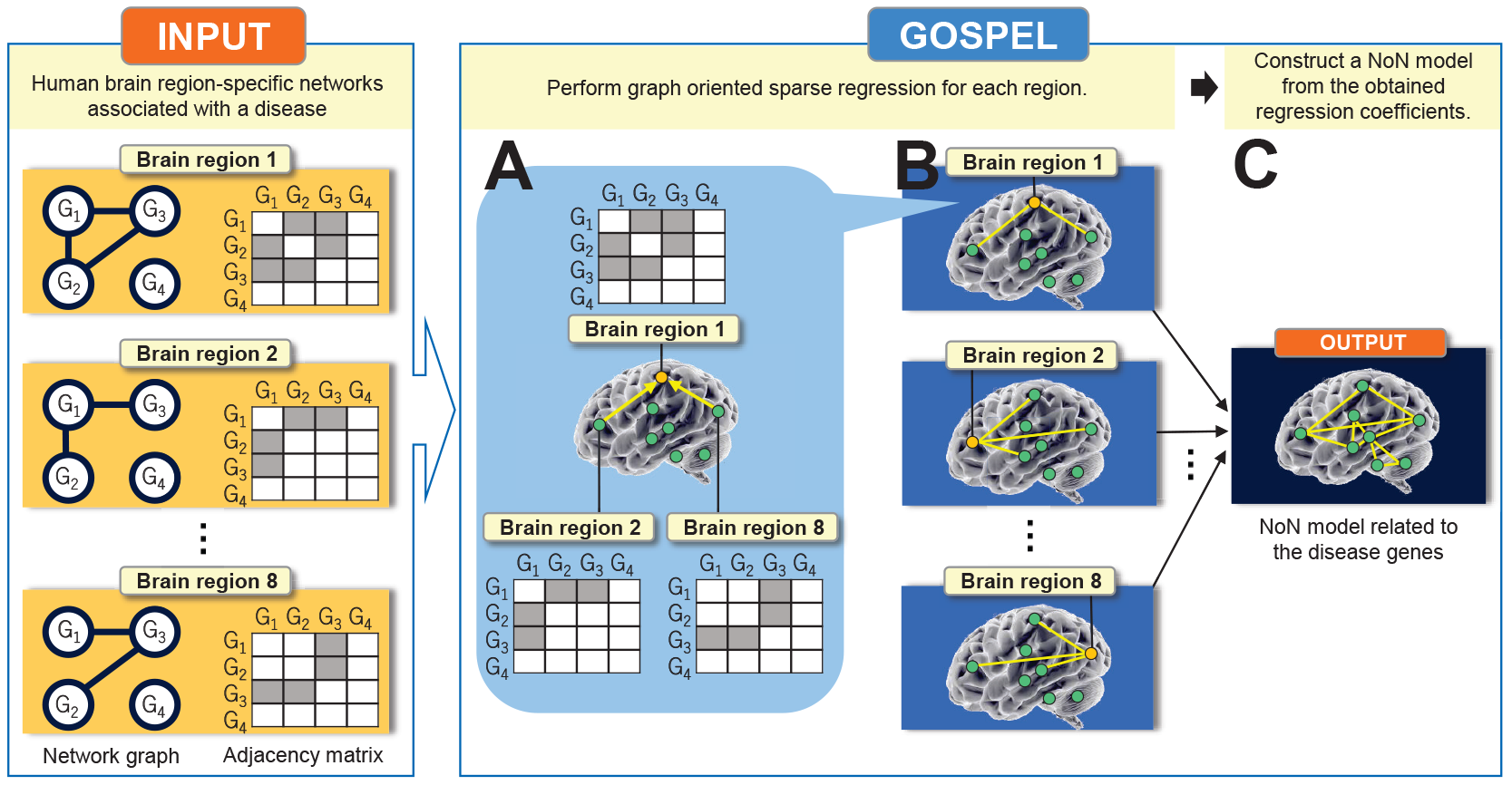
Overview of GOSPEL, an example for the case where *n* = 4, *p* = 8. Assume that we are given the human brain region-specific networks associated with a disease. As input, GOSPEL requires *p* adjacency matrices generated from *p* network graphs with *n* nodes, where *n* and *p* indicate the number of nodes (genes) and features (brainregions) respectively. Block ‘A’ shows that GOSPEL estimates the brain regions which are related to ‘Brain region 1’ by performing a graph oriented sparse regression. In this example, ‘Brain region 2’ and ‘Brain region 8’ are related to ‘Brain region 1’. Block ‘B’ illustrates that aregression is performed on each of the brain regions. As with block ‘A’, block ‘B’ estimates the relationship between the target brain region and the other brain regions. Block ‘C’ expresses that the output of GOSPEL, the “Network of Networks” (NoN) model related to the disease genes, is constructed from the obtained regression coefficients.

For a mechanical understanding of specific genetic variants in neurodevelopmental disorders, integrative network approaches have attracted much attention in recent years due to their interdisciplinary applications. Several state-of-the-art network-based analyses provide an organizational framework of functional genomics and demonstrate that they will enable the investigation of relationships that span multiple levels of analysis (Parikshak *et al.*, 2013; Krishnan *et al.*, 2016; Gandal *et al.*, 2018). These methods were mainly designed for modeling and analyzing single networks that do not interact with or depend on other networks. However, the brain consists of a system of multiple interacting networks and must be treated as such. In multiple interacting networks, the failure of nodes in one network generally leads to the failure of dependent nodes in other networks, which in turn may cause further damage to the first network, leading to cascading failures and catastrophic consequences (Gao *et al.*, 2012). It is known, for example, that different kinds of brain-specific tissues, circuits, and regions are also coupled together and affect behavior and cognitive processes, and thus dysfunctions of the central nervous system in neurodevelopmental disorders have been the result of cascading failures between interdependent systems in the brain. However, no systematic mathematical framework is currently available for adequately modeling and analyzing the consequences of disruptions and failures occurring simultaneously in interdependent networks.

We address this limitation by developing a novel and efficient framework, called the “Network of Networks” (NoN) approach, that will provide useful insights on the properties and topological structure of the inter-correlations between functional interaction networks (Figure. 1). Motivated by a perspective on structural equation models, we model the topological information of each network as a weight sum of the topological information of all other networks. Our NoN model enables the exploitation of the interconnectivity structure between complex systems. It has shown to be effective in aiding the comprehensive understanding of the genetic basis of neurodevelopmental disorders across diverse tissues, circuits, and regions of the brain.

Our main contributions are summarized as follows:

1. We define a statistical framework of structural equation models for inferring the interconnectivity structure between multiple networks where the response and the predictor variables are given networks which have topological information. Structural equation modeling is a statistical method used to test the relationships between observed and latent variables (Civelek, 2018). We extend the structural equation models for modeling the effects of network-network interactions.
2. In order to accomplish this, we propose a sparse learning algorithm for network graph data, called Graph-Oriented SParcE Learning (GOSPEL), to find a subset of the topological information matrices of the predictor variables (networks) related to the response variable (network). More specifically, we propose to use particular forms of diffusion kernel-based *centered kernel alignment* (Cortes *et al.*, 2012) as a measure of statistical correlation between graph Laplacian matrices, and solve the optimization problem with a novel graph-guided generalized fused lasso. This new formulation allows the identification of all types of correlations, including non-monotone and non-linear relationships, between two topological information matrices.
3. We use a Bayesian optimization-based approach to optimize the tuning parameters of the graph-guided generalized fused lasso and automatically find the best fitting NoN model with an acquisition function. The software package that implements the proposed method in the R environment is available from https://github.com/infinite-point/GOSPEL.

We describe our proposed framework and algorithm, and discuss properties in Section 2. Section 3.1 contains a simulation study which demonstrates the performance of the proposed method. We use human brain-specific functional interaction networks and known risk genes with strong prior genetic evidence of ASD and identify the interconnectivity between these networks in Section 3.2. Section 4 provides concluding remarks.

## 2 Method

Our goal is to infer the interconnectivity structure between multiple networks from topological information matrices of given networks. To do this, we make the assumption that each topological information matrix for a given network can be expressed by the linear combination of the topological information matrices of the other given networks. Sparse regression is performed on each of the given networks in order to identify a subset of the topological information matrices of the predictors related to the response. After this computation, NoN model is constructed from the obtained regression coefficients. In this section, we first explain the problem setting, and then present our method, GOSPEL.

### 2.1 Problem Setting

Suppose that we are given *p* undirected network graphs consisting of *n* vertices (nodes) 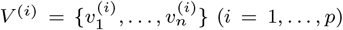 linked by edges. The *i*-th *adjacency* matrix ***A***^(*i*)^ ∈ ℝ^*n*×*n*^ associated with the *i*-th undirected network graph is defined as

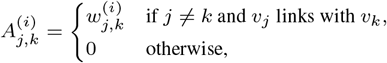

where 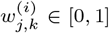 denotes the probability of connectivity between the *v_j_* and *v_k_* in the *i*-th network graph. Here, we compute *graph Laplacian* matrix ***L***^(*i*)^:

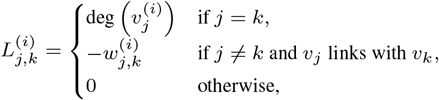

where deg 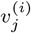 denotes the degree of 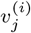. Then let us kernelize the *graph Laplacian* matrix. Let ***K***^(*i*)^ be the *diffusion kernel* matrix for ***L***^(*i*)^:

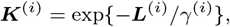

where *γ^(i)^* is a kernel parameter. This kernel matrix is centered and normalized as follows:

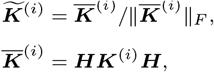

where ‖ · ‖*_F_* denotes the Frobenius norm, ***H*** indicates the centering matrix 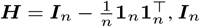, ***I***_*n*_ is an *n* × *n* identity matrix and **1**_*n*_ is an *n*-dimensional vector with all ones.

We assume that the diffusion kernel matrix of the *i*-th network graph can be represented by the linear combinations of the diffusion kernel matrices of the other network graphs as follows:

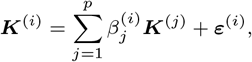

where 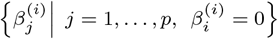 denotes a regression coefficient corresponding to predictor ***K***^(*j*)^ and response ***K***^(*i*)^, and *ε*^(*i*)^ ∈ ℝ^*n*×*n*^ is a Gaussian noise matrix whose elements follow 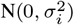.

The estimation for the network structure of the given networks is available by applying GOSPEL to all the cases where *i* = 1,…,*p*. Note that, in an ordinary regression problem for *n* samples and *p* predictors, both the predictors and the response are given as *n*-dimensional vectors. In our problem setting, however, they are given as *n* × *n* network graphs.

**Table 1.**
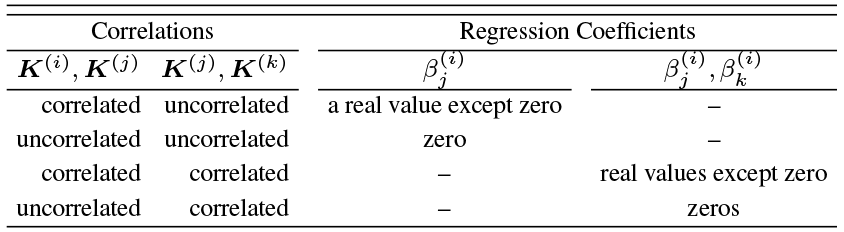
Behaviour of 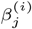 and 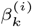 in GOSPEL optimization. When ***K***^(*j*)^ and ***K***^(*k*)^ are uncorrelated, i.e.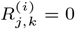 the value of 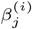 is estimated depending on the correlation between response ***K***^(*i*)^ and predictor ***K***^(*j*)^. Similarly, the value of 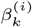 is computed depending on the correlation between the responseandthe *k*-thpredictor.Ontheotherhand,when ***K***^(*j*)^ and ***K***^(*k*)^ are correlated, i.e.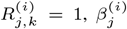 and 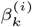 tend to take similar values depending on the correlation between the response and the *i*-th predictor.

### 2.2 Graph Oriented Sparse Learning (GOSPEL)

The optimization problem of GOSPEL is as follows:

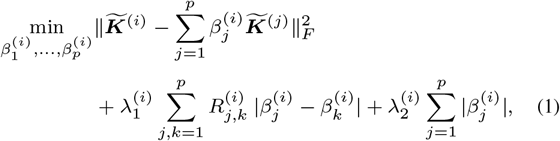

where 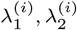 are regularization parameters and | · | indicates the *ℓ*_1_ norm. ***R***^(*i*)^ ∈ ℝ^*p* × *p*^ (*i* = 1,…,*p*) expresses a matrix whose elements consist of correlations between predictors, where

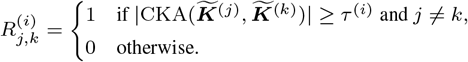

CKA(***K***^(*j*)^, ***K***^(*k*)^) denotes a correlation between kernel matrices ***K***^(*j*)^ and ***K***^(*k*)^; this measure is called the *Centred Kernel Alignment* (**CKA**) (Cortes *et al.*, 2012), and τ^(*i*)^ indicates a threshold. CKA captures the non-linear relationship between two matrices if such a relationship exists. The definition of CKA is as follows:
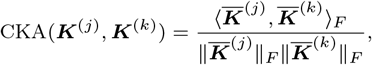

where 〈·,·〉_*F*_ indicates the Frobenius inner product. The Frobenius inner product can be interpreted as an inner product of two vectorized matrices, and thus we can apply the properties of Pearson’s correlation coefficient (Sharma, 2005) to CKA. Unless the elements of ***K***^(*j*)^ or ***K***^(*k*)^ are all zero (we omit such cases in the computation of GOSPEL), this definition implies that the value of CKA becomes zero when ***K***^(*j*)^ and ***K***^(*k*)^ have no correlation and the CKA value takes ±1 when the two matrices are strongly correlated. In practice, the value of the *diffusion kernel* based CKA ranges from –1 to 1 because of the positive semi-definiteness of the *diffusion kernel* matrix (Lafferty and Kondor, 2002).

GOSPEL optimization Eq. (1) consists of the squared Frobenius norm term, the *graph-guided-fused-lasso* regularization term (Chen *et al.*, 2012) and the *lasso* regularization term (Tibshirani, 1996). If 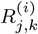 of the *graph-guided-fused-lasso* regularization term is zero, the equation becomes solely dependant on the *lasso* regularization term. Table 1 summarizes the behavior of 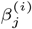 and 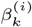 in GOSPEL optimization. The table shows that GOSPEL estimates all the relevant predictor networks to the response network, and also eliminates irrelevant predictor networks to the response network. For more detail on the behavior of the regression coefficients, see section 1 in the supplement. The sparsity of the elements of *β*^(*i*)^ helps to facilitate the interpretation of the computation results. By extension, the interpretation of the network structure of given networks is also facilitated.

Finally, we construct a NoN model utilizing the obtained regression
coefficients 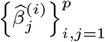. Let *E* ⊆ Γ(*p*) × Γ(*p*) be an edge set for a NoN model, where Γ = {1,…,*p*} is a node set. We employ the edge set estimation defined by Meinshausen and Bühlmann (2006):

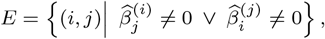

where (*i, j*) indicates the pair of the *i*-th and the *j*-th nodes. Based on the graphical model *G* = (Γ, *E*), the NoN model is constructed as the output of GOSPEL.

### 2.3 Computation of GOSPEL

To solve the GOSPEL optimization problem, Eq. (1), we first vectorize all the kernel matrices. This produces an *n*^2^-dimensional vector associated with the response network, and *n*^2^-dimensional vectors corresponding to *p* − 1 predictor networks. This form is the same problem setting of the *graph guided, generalized fused lasso* (Chen *et al.*, 2012), G3FL, with *n*^2^ samples and *p* features. Therefore, we employ G3FL to solve our optimization problem.

Regularization parameters 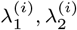, threshold τ^(*i*)^ and kernel parameter γ^(*i*)^ are decided by the *Bayesian Optimization* (Mockus, 2012). We apply the Bayesian Information Criterion (BIC) (Schwarz, 1978) as an *acquisition function* of the *Bayesian Optimization.* The BIC score for the case where the response is the *i*-th network graph is defined as

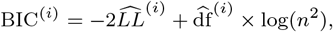

where 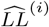 is the log-likelihood function:

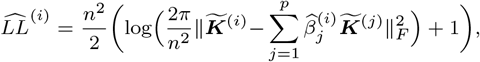

and 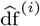 is the degree of freedom of the *fused lasso* (Tibshirani *et al.*,2005):

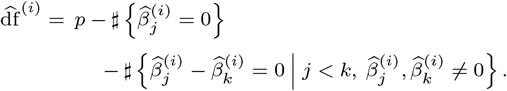

In the *Bayesian Optimization*, we select the set of parameter values which minimizes the BIC score.

## 3 Results

### 3.1 Simulations

We generate synthetic data and evaluate the performance of GOSPEL in order to gain insight into feature selection in the regression problem for network graph data. As synthetic data, we prepare three representative complex network models which have different structures: random networks (Erdös and Rényi, 1959), scale-free networks (Barabási and Albert, 1999) and small-world networks (Watts and Strogatz, 1998). For each network model, 30 predictor networks are prepared so that the first 15 predictors have the following non-linear relationships:

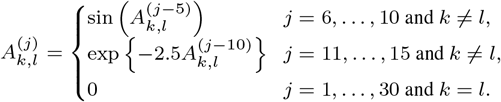

Using the 15 predictors, we generate the following two signal types of response networks with signal-to-noise ratio equal to 1.
- Additive type: 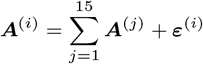
- Non-additive type: 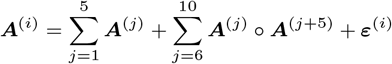
where ○ indicates the element-wise product.

We run the simulations 100 times for each combination of the network model and the signal type, varying the number of vertices as *n* = {500, 1000, 2000}. We compare GOSPEL to HSIC Lasso (Yamada*etal.*, 2014), one of the feature selection methods by the feature-wise kernelized lasso. We note that HSIC Lasso is not designed for network graph data and cannot be directly applied. In these simulations, HSIC lasso is applied to the centered and normalized kernel matrices as follows:

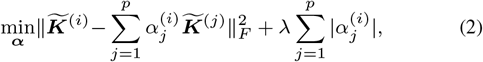

where 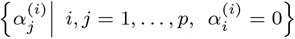

To assess each method’s ability to obtain true fractions of predictors related with the response, we compare true predictors to the predictors with non-zero coefficients estimated by GOSPEL and HSIC Lasso. The results were analyzed for precision, recall and F-measure. Table 2 shows the F-measures calculated for the 18 different settings with varying sample size and network types. Regarding the precision and the recall of the simulation results, see Tables 1 and 2 in the supplement, respectively. The results highlight the efficacy of the *graph-guided fused-lasso* regularization.

In the cases of random networks, GOSPEL’s estimation performance remains high regardless of the sample size or the signal type. This result may come from the fact that the random network is the simplest network of the three. Compared with the random network, the scale-free and small-world networks are difficult to estimate. Since the scale-free and small-world networks have distinctive structures, all the predictor networks are similar to each other within their group. In the case of the scale-free network, when the signal type is additive, the performance of GOSPEL becomes better as *n* grows. On the other hand, when the signal type is non-additive, GOSPEL and the variant of HSIC Lasso perform almost at the same level. The reason for this is that most elements of the response network take similar values due to the element-wise product
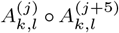. Since the operation tends to break the structure of the response network in spite of the predictor networks keeping their structures, estimation becomes excessively difficult. Interestingly, the performance of the non-additive cases is better than that of the additive cases in the simulations of small-world networks. In the non-additive cases,the operation of the element-wise product may work to emphasize the structure of the response network, and this may improve performance here while leading to opposite results in the scale-free cases.

Our simulation results demonstrate that GOSPEL is able to recover the true network structure from given network graph data, and outperforms HSIC Lasso in terms of precision, recall and F-measure in the different settings with varying sample size and network types.

### 3.2 Real data

ASD is a complex neurodevelopmental disorder driven by a multitude of genetic variants across the genome that appear as a range of developmental and functional perturbations, often in specific tissues and cell types (Vorstman *etal.*, 2017). To construct human brain region-specific networks associated with the ASD risk genes, we adopt a manner of data-construction introduced in recent studies (Krishnan *et al.*, 2016; Duda *et al.*, 2018). We use 17 human brain region-specific functional interaction networks (Greene *et al.*, 2015) and 1030 known risk genes with strong genetic evidence of ASD association annotated in the Human Gene Mutation Database (HGMD) Professional 2017.1 (http://hgmd.cf.ac.uk/) to evaluate our proposed method. The purpose of our analysis is to investigate how the ASD risk genes may be coupled in each of the brain region-specific networks and what inter-connectivity structure between these networks can be formed.

**Table 2.** The F-measure of the simulations. (·) denotes the standard deviation of the F-measure. The predictor networks are generated according to the representative three network models. *n* indicates the number of vertices, and the response network is generated based on the signal type. ‘HSIC Lasso’ does not mean the original one but the variant one.

| $n$ | Signal type | Network Model | | | | | |
| --- | --- | --- | --- | --- | --- | --- | --- |
|  |  | Random |  | Scale-free |  | Small-world |  |
|  |  | GOSPEL | HSIC Lasso | GOSPEL | HSIC Lasso | GOSPEL | HSIC Lasso |
| 500 | Additive | 0.987 | 0.737 | 0.883 | 0.516 | 0.893 | 0.789 |
|  |  | (0.017) | (0.069) | (0.085) | (0.024) | (0.078) | (0.039) |
|  | Non-additive | 0.986 | 0.589 | 0.674 | 0.639 | 0.927 | 0.688 |
|  |  | (0.024) | (0.034) | (0.015) | (0.023) | (0.047) | (0.045) |
| 1000 | Additive | 0.985 | 0.746 | 0.894 | 0.541 | 0.973 | 0.703 |
|  |  | (0.019) | (0.058) | (0.145) | (0.032) | (0.022) | (0.061) |
|  | Non-additive | 0.988 | 0.588 | 0.678 | 0.604 | 0.988 | 0.540 |
|  |  | (0.029) | (0.038) | (0.020) | (0.028) | (0.024) | (0.023) |
| 2000 | Additive | 0.985 | 0.787 | 0.932 | 0.510 | 0.900 | 0.800 |
|  |  | (0.018) | (0.054) | (0.135) | (0.016) | (0.057) | (0.006) |
|  | Non-additive | 0.995 | 0.592 | 0.688 | 0.635 | 0.962 | 0.721 |
|  |  | (0.014) | (0.038) | (0.036) | (0.015) | (0.031) | (0.056) |

The 17 human brain regions are taken from the whole brain: the frontal lobe (the cerebral cortex), the parietal lobe (the cerebral cortex), the temporal lobe (the cerebral cortex), the occipital lobe (the cerebral cortex), the subthalamic nucleus (the basal ganglia), the caudate nucleus (the basal ganglia), the caudate putamen (the basal ganglia), the amygdala (the limbic system), the nucleus accumbens (the limbic system), the hippocampus (the limbic system), the dentate gyrus (the limbic system), the hypothalamus (the diencephalon), the thalamus (the diencephalon), the hypophysis (the diencephalon), the substantia nigra (the brain stem), the pons (the brain stem), and the cerebellar cortex (the cerebellum).

These 17 human brain region-specific functional interaction networks are built by integrating thousands of gene expressions, protein-protein interactions, and regulatory-sequence data sets using a regularized Bayesian integration approach (Greene *et al.*, 2015). Once built, they are used as reference networks in our analysis. This network information can be downloaded from the Genome-scale Integrated Analysis of gene Networks in Tissues (GIANT) web site (http://giant.xsprinceton.edu/).

We first calculate local enrichment scores for the ASD risk genes across all nodes (25825 genes) in the network by using the Spatial Analysis of Functional Enrichment (SAFE) algorithm (Cerebral, 2016), which measures the proximity of the ASD risk genes in the neighborhood of each node. Next, in order to reconstruct the 17 human brain region-specific networks associated with the ASD risk genes, 4756 genes with highly significant local enrichment scores for the ASD risk genes (p-value < 0.001) in any of the 17 networks are selected as the nodes, and the interactions between these genes are given. We apply GOSPEL to the 4756 × 4756 adjacency matrices of the 17 brain-specific regions and construct the NoN model related to the ASD risk genes.

Figure. 2 shows a NoN model related to the ASD risk genes. It illustrates the relation between the functional networks of the subregions. This result suggests the existence of the topological structure of the inter-correlation between the functional interaction networks associated with ASD. For the interpretation of the resulting model, we perform a community extraction method based on random walks (Pons and Latapy, 2006). Table 3 indicates the mean value of enrichment score and the community ID for each subregion, and shows that the resulting model is divided into 3 communities. The first group is characterized by the amygdala and the thalamus. The amygdala has the largest number of the thickest edges and the thalamus is the hub in this group. The second group is small, however, it has the interesting feature that two out of the three subregions belong to cerebral cortex. The third group consists of the subregions which have low enrichment sores, and thus we consider this group as not relating to ASD and ignore it here.

**Table 3.** The mean value of enrichment score and the community ID for each subregion. Enrich.score and Com.ID indicate the mean value of enrichment score and the community ID, respectively.

| Subregion | Enrich.score | Com.ID |
| --- | --- | --- |
| Temporal lobe | 10.322 | 1 |
| Hippocampus | 10.320 | 1 |
| Hypophysis | 10.261 | 1 |
| Caudate nucleus | 9.890 | 1 |
| Thalamus | 9.386 | 1 |
| Amygdala | 9.259 | 1 |
| Substantia nigra | 7.967 | 1 |
| Subthalamic nucleus | 7.955 | 1 |
| Frontal lobe | 8.130 | 2 |
| Occipital lobe | 7.212 | 2 |
| Caudate putamen | 5.476 | 2 |
| Cerebellar cortex | 5.402 | 3 |
| Dentate gyrus | 3.879 | 3 |
| Hypothalamus | 3.485 | 3 |
| Nucleus accumbens | 3.290 | 3 |
| Pons | 2.849 | 3 |
| Parietal lobe | 1.318 | 3 |

Table 4 shows the consistency between our resulting model and the abnormal functional connections in Table 1 of Yahata *et al.* (2016), and samples of the evidence associated with each subregion and ASD. See Table 3 in the supplement for further information. For the classification of ASD and typically developed (TD) persons, Yahata *et al.* (2016) identified the 16 abnormal functional connections using fMRI. We investigate the consistency of the subregions studied in Yahata *et al.* (2016) and those in our experiment. There are five connections corresponding to our experiment: the caudate nucleus and the amygdala, the frontal lobe and the occipital lobe, the temporal lobe and the frontal lobe, the hippocampus and the frontal lobe, and the frontal lobe and the parietal lobe. As shown in Table 4, four out of the five connections are shared.

In the first group, the amygdala has the largest number of the thickest edges, and thus the amygdala is suggested to be the center of functional abnormality in this group, whereas the thalamus acts as the hub. Table 4 supports this suggestion; the amygdala is not only included in the abnormal functional connection in Yahata *et al.* (2016) but is also well studied as a subregion strongly related to ASD. As stated, the hub of the first group is the thalamus, and this result is analogous to the medical knowledge that the thalamus is an information relay station (hub) between the subcortical areas and the cerebral cortex (Gazzaniga *et al.*, 2009). In addition, some links in the first group, such as the hippocampus and the temporal lobe, and the caudate nucleus and the thalamus, are physically close even though we did not consider locational information in this experiment.

The subregion which seems to be representative of the second group is the frontal lobe. As shown in Table 4, the frontal lobe is a well-studied subregion in ASD research. In addition, four out of the five abnormal connections corresponding to our experiment shown above include the frontal lobe. It is remarkable that the resulting model shares three out of the four abnormal connections.

**Fig. 2.**
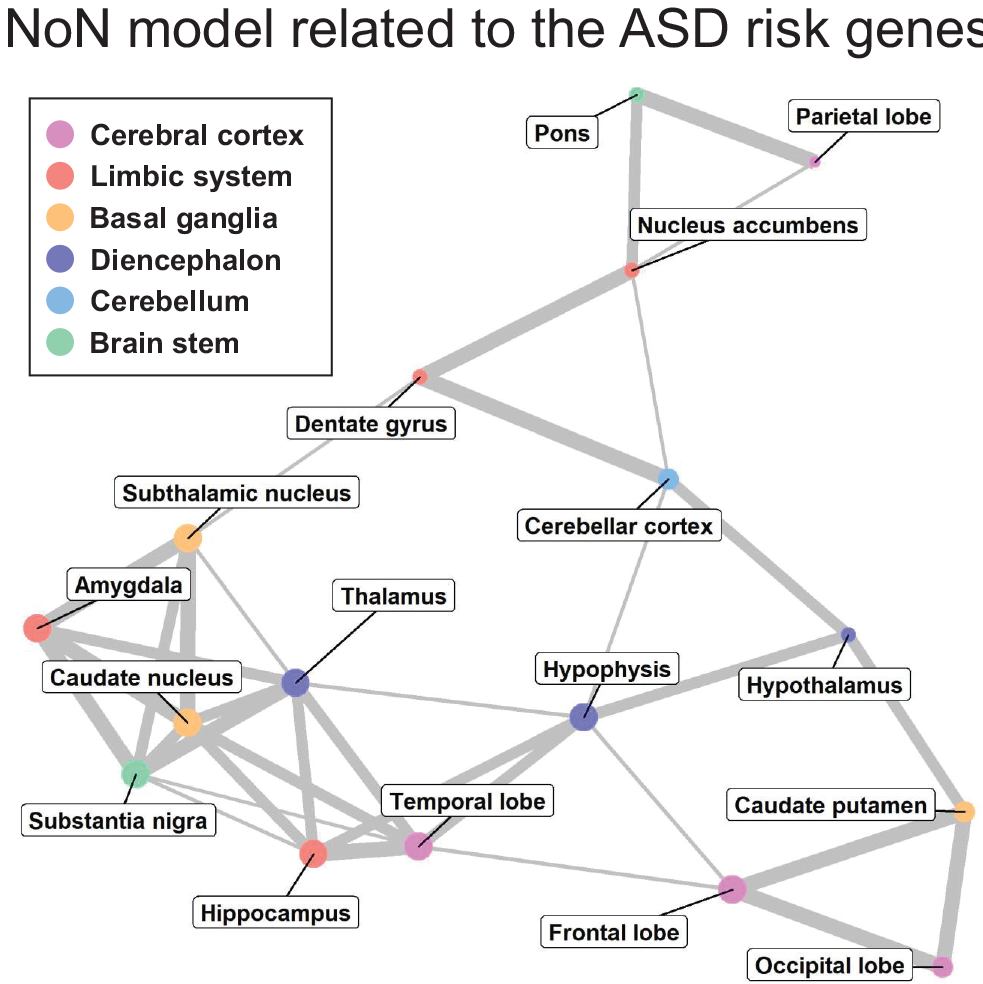
NoN model related to the ASD risk genes. The size of the node expresses the mean value of the enrichment score associated with the brain subregion (node). The smallest, middle and the largest sized nodes express below the 70-th percentile, between the 70-th and the 85-th percentile and above the 85-th percentile of the enrichment score, respectively. The color of the node indicates the anatomical classification. The thickness of the edge denotes the strength of the relation between two brain subregions. The thinnest, middle and the thickest egdes express the 70-th to the 80-th percentile, the 80-th to the 90-th percentile and above the 90-th percentile of the whole coefficients obtained by performing GOSPEL, respectively. For details on edge thickness (edge weight), see Table 4 in the supplement.

**Table 4.** Consistency between our resulting model and the abnormal functional connections in Table 1 of Yahata *et al.* (2016), and samples of the evidence for a single brain subregion associated with ASD. ‘◯’ and ‘△’ denote a direct and a proximate connection, respectively.

| Com.ID | Subregion | Consistency | Samples of the evidence for a single brain subregion |
| --- | --- | --- | --- |
| 1 | Temporal lobe | ○ | Zilbovicius *et al.* (2000); Lee *et al.* (2007); Neeley *et al.* (2007). |
|  | Hippocampus | △ | Schumann *et al.* (2004); Endo *et al.* (2007); Conturo *et al.* (2008). |
|  | Hypophysis |  | Iwata *et al.* (2011); Ćurin *et al.* (2003). |
|  | Caudate nucleus | ○ | Silk *et al.* (2006); Turner *et al.* (2006); Voelbel *et al.* (2006); Langen *et al.* (2007). |
|  | Thalamus |  | Tsatsanis *et al.* (2003); Hardan *et al.* (2006); Hardan *et al.* (2008); Tamura *et al.* (2010). |
|  | Amygdala | ○ | Schumann *et al.* (2004); Schumann and Amaral (2006); Endo *et al.* (2007); Conturo *et al.* (2008); Schumann *et al.* (2009); Kleinhans *et al.* (2010); Nordahl *et al.* (2012); Kliemann *et al.* (2012). |
|  | Substantia nigra |  | Caria *et al.* (2011). |
|  | Subthalamic nucleus |  | Rojas *et al.* (2014). |
| 2 | Frontal lobe | ○ ○ △ | Hardan *et al.* (2004); Schmitz *et al.* (2007); Scott-Van Zeeland *et al.* (2010); Jeong *et al.* (2011). |
|  | Occipital lobe | ○ | Hardan *et al.* (2009). |
|  | Caudate putamen |  | Kumra *et al.* (2000). |

The analysis with a real example thus shows that GOSPEL is able to identify the ASD-associated interconnectivity structure between given functional interaction networks.

## 4 Discussion

In order to improve understanding of brain-specific complex systems related to a disease and to break through the limitation of the network-based analyses which estimate functional single networks, we proposed a “Network of Networks” (NoN) approach inspired by structural equation models. In this paper, we sought to estimate the topological structure of the correlations between functional interaction networks of human brain region-specific networks associated with ASD risk genes.

To the best of our knowledge, the sparse regression for network graphs in GOSPEL is the first feature selection method where the features are not vectors but instead are network graphs. In order to construct a NoN model, GOSPEL estimates all the predictor networks relevant to the response network even when they have non-linear correlations. All the parameters in GOSPEL are automatically optimized by the *Bayesian Optimization* based on the BIC. Though there is room for improvement in that GOSPEL can not reflect the information of the vertices (nodes), the outputted NoN model is interpretable by combining the information on the vertices and the result of community extraction, as shown in Section 3.2.

We demonstrated the effectiveness of GOSPEL in simulations, and tackled the exploration of the NoN model of human brain region-specific networks related to ASD. The result was the successful production of a NoN model which shows the subregions of the brain which relate to ASD and how they functionally relate to each other. This model is consistent with previous ASD research. Although the accuracy of our model is yet untested, we hope that it will provide useful insight for ASD researchers, and that further research will prove its accuracy.

Finally, while this research was limited to the study of subregions of the brain, we believe that GOSPEL will prove useful to other studies seeking to find relationships between illness and bodily organs or regions, therefore it may be of great use to those studying the *systems of biology.*

## Acknowledgements

We would like to thank Makoto Yamada for helpful discussions.

